# Extended functional connectivity of convergent structural alterations among individuals with PTSD: A neuroimaging meta-analysis

**DOI:** 10.1101/2022.04.07.487478

**Authors:** Brianna S. Pankey, Michael C. Riedel, Isis Cowan, Jessica E. Bartley, Rosario Pintos Lobo, Lauren D. Hill-Bowen, Taylor Salo, Erica D. Musser, Matthew T. Sutherland, Angela R. Laird

## Abstract

**Background:** Post-traumatic stress disorder (PTSD) is a debilitating disorder defined by the onset of intrusive, avoidant, negative cognitive or affective, and/or hyperarousal symptoms after witnessing or experiencing a traumatic event. Previous voxel-based morphometry studies have provided insight into structural brain alterations associated with PTSD with notable heterogeneity across these studies. Furthermore, how structural alterations may be associated with brain function, as measured by task-free and task-based functional connectivity, remains to be elucidated.

**Methods:** Using emergent metaanalytic techniques, we sought to first identify a consensus of structural alterations in PTSD using the anatomical likelihood estimation (ALE) approach. Next, we generated functional profiles of identified convergent structural regions utilizing resting-state functional connectivity (rsFC) and meta-analytic coactivation modeling (MACM) methods. Finally, we performed functional decoding to examine mental functions associated with our ALE, rsFC, and MACM brain characterizations.

**Results:** We observed convergent structural alterations in a single region located in the medial prefrontal cortex. The resultant rsFC and MACM maps identified functional connectivity across a widespread, whole-brain network that included frontoparietal and limbic regions. Functional decoding revealed overlapping associations with attention, memory, and emotion processes.

**Conclusions:** Consensus-based functional connectivity was observed in regions of the default mode, salience, and central executive networks, which play a role in the tripartite model of psychopathology. Taken together, these findings have important implications in understanding the neurobiological mechanisms associated with PTSD.

## Background

Post-traumatic stress disorder (PTSD) is a psychiatric disorder in which the onset of symptoms develops after experiencing or witnessing a traumatic event, such as violence, accidents, or combat (Yehuda et al., 2015). Symptoms associated with PTSD are categorized into clusters according to the DSM 5: (1) intrusion/re-experiencing trauma, (2) avoidance, (3) negative cognition and mood, and (4) hyperarousal (Kirkpatrick and Heller, 2014; Pai et al., 2017). Approximately 70% of adults experience at least one traumatic event in their lifetime and up to 20% of these people develop PTSD (PTSD Alliance, 2018). Individuals with PTSD may experience long-term debilitating effects, mentally, physically, and cognitively. In the United States, roughly 8 million adults suffer from PTSD every year. Approximately 60% of men experience at least one traumatic event in their lives, often associated with combat and war, while 50% of women will experience at least one traumatic event, typically associated with sexual assault and abuse (National Center for PTSD, 2019).

Current theories aim to understand the etiology of PTSD, including behavioral, cognitive, and social models. Research suggests that reappraisal of traumatic events may lead to an overgeneralized threat response (Ehlers and Clark, 2000). Despite progress in understanding the vulnerability, symptomatology, and trajectory of PTSD (Agaibi and Wilson, 2005; Pitman et al., 2012; Kirkpatrick and Heller, 2014), the underlying neurobiological determinants of PTSD are less clear. Substantial prior work has attempted to identify structural brain alterations observed among individuals with PTSD. Voxelbased morphometry (VBM) is a commonly used methodological approach for analyzing structural magnetic resonance imaging (MRI) data, allowing for quantitative statistical comparisons between groups (e.g., differences in gray matter volume; GMV) to more clearly understand structural alterations linked to neuropsychiatric disorders, such as PTSD. Multiple prior meta-analyses have been conducted to identify convergent gray matter reductions in PTSD patients, although consensus *across meta-analyses* has not been reached. Each of these meta-analyses was conducted with a different scope, with varied study inclusion/exclusion criteria, and subsequently included a wide range of 8 to 20 studies. Varying convergence has been observed across these meta-analyses, which have identified one to five significant clusters in regions that include medial prefrontal cortex (Kühn and Gallinat, 2013; Li et al., 2014; Meng et al., 2014; Bromis et al., 2018; Klaming et al., 2019), hippocampus (Kühn and Gallinat, 2013; Bromis et al., 2018), fusiform gyrus (Li et al., 2014; Serra-Blasco et al., 2021), and lingual gyrus (Kühn and Gallinat, 2013; Serra-Blasco et al., 2021). Similarly, from a functional perspective, PTSD dysfunction has been reported as amygdala and frontal disruptions (e.g., Duval et al., 2015) or across alterations of large-scale functional brain networks (e.g., Koch et al., 2016) that are implicated in the tripartite model of psychopathology (Menon, 2011). While some studies have addressed consensus across functional neuroimaging studies, it is challenging to assess convergence across different psychological states and/or experimental paradigms, which has potentially contributed to inconsistent findings in PTSD meta-analyses of resting state (Wang et al, 2016; Bao et al., 2021) or task-based (Etkin and Wager, 2007; Patel et al., 2012; Hayes et al., 2016) studies. Overall, this variability across meta-analytic approaches and results suggests that a consensus neurobiological model of PTSD has not yet been achieved.

The objective of the current study was to apply current best practices in coordinate-based neuroimaging methods to investigate the topography of consistently reported structural alterations in PTSD. As PTSD is linked to a broad spectrum of neuropsychiatric symptoms, which likely reflects the disturbance of distributed, brain-wide neural circuitry, we also sought to functionally and behaviorally characterize any neuroanatomical alterations in a task-independent manner. To this end, we first identified convergent regions of gray matter (GM) reductions in PTSD vs. non-PTSD groups using anatomical likelihood estimation (ALE) (Eickhoff et al., 2009, 2012). Second, we identified the task-free resting state functional connectivity (rsFC) patterns, as well as the task-based meta-analytic co-activation modeling (MACM) patterns of convergent regions, thus providing multimodal functional connectivity profiles for each. Together, the VBM, rsFC, and MACM meta-analytic approaches have been used in previous clinically related meta-analyses (Reetz et al., 2012; Dogan et al., 2015; Kamalian et al., 2022); they provide complementary information, yielding a multimodal functional connectivity profile for a given region of interest. Lastly, we applied meta-analytic functional decoding methods to identify the mental processes linked to this functional connectivity profile. Collectively, this work utilizes an innovative (meta-)analytic framework to quantitatively assess structural alterations associated with PTSD and the extended functional profiles of regions implicated in this disorder. A more comprehensive understanding of the neurobiological bases of PTSD is needed to delineate future pathways toward improved prevention, diagnosis, and treatment.

## Methods

### Analytic Overview

We first conducted a literature search to identify studies reporting structural alterations comparing the following groups: individuals with PTSD, individuals who experienced trauma but were not diagnosed with PTSD, and individuals who did not report experiencing trauma. A coordinate-based meta-analysis was performed using the ALE algorithm to identify convergent brain regions showing structural alterations associated with PTSD. We then used multiple connectivity modeling approaches to comprehensively characterize the functional connectivity of these convergent regions. Specifically, rsFC and MACM assessments were applied to identify the functional profiles of structurally altered regions associated with PTSD. Lastly, we used functional decoding techniques to identify behavioral profiles of the ALE, rsFC, and MACM results. An overview of our methodological approach is provided in **Figure 1**.

**Figure 1.**
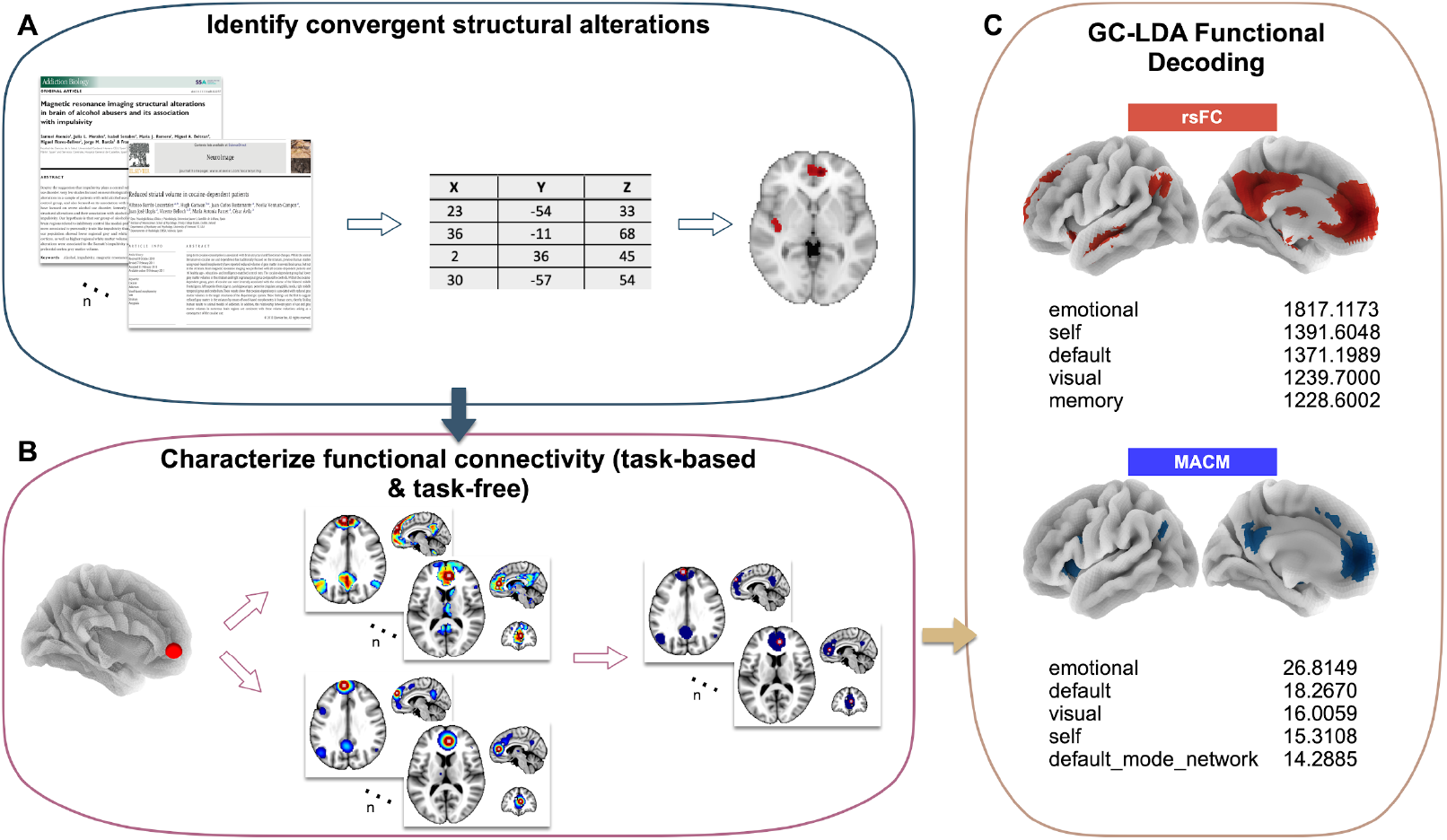
Analysis Pipeline Overview. A) We first conducted a literature search to extract structural coordinates and entered them into the ALE algorithm to identify convergent structural alterations among PTSD vs. non-PTSD groups. B) We next created task-free and task-based functional connectivity profiles for the convergent structural alterations. C) Last, we performed functional decoding analyses on these functional profiles to make inferences about which mental functions were associated with our findings.

### Literature Search and Study Criteria

We conducted a comprehensive literature search to build a database of peer-reviewed MRI studies reporting structural alterations associated with PTSD from 2002-2020. In the first round of identifying studies, we examined previously published voxel-based morphometry meta-analysis papers on PTSD and compiled a list of included studies (Kühn and Gallinat, 2013; Li et al., 2014; Meng et al., 2014; Bromis et al., 2018; Klaming et al., 2019). Next, we performed a PubMed search to identify additional peer-reviewed, structural MRI studies of interest using the search terms “morphometry + PTSD”. The PubMed search aimed to identify any potential studies that were not included in the previously published meta-analyses. We then conducted a review of each identified publication to include the following study criteria: peer-reviewed MRI studies, reporting results among adult humans, written in the English language, focused on gray matter structural differences, and included original data (i.e., not a review). Subsequently, exclusion criteria were as follows: trauma or stressful life event studies not measuring PTSD, other non-voxel-based morphometry methods, treatment and longitudinal effects, papers reporting *a priori* regions of interest (ROIs), within-group effects, null effects, overlapping samples to previous studies, and studies that did not report coordinate-based results.

### Anatomical Likelihood Estimation (ALE)

ALE is a voxel-based meta-analytic technique that identifies convergent coordinates (i.e., foci) across a set of neuroimaging studies. Foci are treated as 3D Gaussian distributions to address variability within and between studies. We used the coordinate-based ALE method as implemented in NiMARE v.0.0.3 (*Neuroimaging Meta-Analysis Research Environment*; Salo et al., 2022), a Python library for neuroimaging meta-analysis. Reported coordinates were extracted from their original publication; coordinates originally reported in Talairach space converted to were MNI coordinates (Lancaster et al., 2007; Laird et al., 2010) so that all coordinates referred to MNI space. Once transformed, statistical probability maps were created for each foci and combined to model the likelihood that a given voxel displayed a between-group structural difference for each study. Observed voxel-wise ALE scores characterized the most consistently reported foci across the whole brain. Significance testing and correction for multiple comparisons involved thresholding the voxel-wise ALE map using a cluster-forming threshold of *P* < 0.001. Then, a permutation procedure was performed in which a null distribution of maximum cluster sizes was generated from 10,000 iterations of replacing reported foci with randomly selected gray matter voxels, generating ALE maps from the randomized dataset, and identifying the maximum cluster size after thresholding at *P* < 0.001. The cluster-level FWE correction threshold was set at *P* < 0.05, meaning only those clusters from the original, thresholded ALE map were retained if their size was greater than the cluster size corresponding to the 95th-percentile from the null distribution. We applied the above ALE procedure to identify convergent brain regions reflecting structural alterations between individuals with and without PTSD (i.e., PTSD vs. non-PTSD) separately for the contrasts of PTSD > non-PTSD and non-PTSD > PTSD.

### Functional Profiles of Structurally Altered Regions Associated with PTSD

Next, we sought to characterize the functional connectivity patterns associated with regions demonstrating structural alterations in PTSD. To this end, we investigated *task-free functional connectivity* utilizing a database of resting state fMRI data, as well as *task-based functional connectivity* using a meta-analytic database of co-activation results.

### Task-Free Functional Connectivity: Resting-State fMRI (rs-fMRI)

Resting-state connectivity analyses typically identify brain voxels demonstrating the highest temporal correlation with the average time series of a seed ROI and provide context about the brain’s underlying functional architecture. To derive robust rsFC maps for each ROI, we utilized the minimally pre-processed and denoised (or “cleaned”) resting-state fMRI data provided by the Human Connectome Project’s (Van Essen et al., 2013) Young Adult Study S1200 Data Release (March 1, 2017). On November 12, 2019, 150 randomly selected participants (28.7 ± 3.9 years) were downloaded via the HCP’s Amazon Web Services (AWS) Simple Storage Solution (S3) repository. The randomly chosen participants included 77 females (30.3 ± 3.5 years) and 73 males (27.1 ± 3.7 years). A difference in age between the two biological sex groups was significant but is consistent with the 1200 Subjects Data Release. Detailed acquisition and scanning parameters for HCP data can be found in consortium manuscripts (Van Essen et al., 2012; Smith et al., 2013; Uğurbil et al., 2013), but relevant scan parameters are briefly summarized here. Each participant underwent T1weighted and T2-weighted structural acquisitions and four resting-state fMRI acquisitions. Structural images were collected at 0.7-mm isotropic resolution. Whole-brain EPI acquisitions were acquired on the 3T Siemens Connectome scanner: 32-channel head coil, TR = 720 msec, TE = 33.1 msec, in-plane FOV = 208 × 180 mm, 72 slices, 2.0 mm isotropic voxels, and multiband acceleration factor of 8 (Feinberg et al., 2010).

The S1200 data release contained minimally pre-processed and denoised data. The minimal preprocessing workflow is described by Glasser and colleagues (Glasser et al., 2016), but consists of typical imaging pre-processing techniques that leverage the high-quality data acquired by the HCP. First, T1- and T2-weighted images were aligned, bias field corrected, and registered to MNI space. Second, the functional fMRI pipeline removed spatial distortions, realigned volumes to compensate for subject motion, registered the fMRI data to structural volumes (in MNI space), reduced the bias field, normalized each functional acquisition to its corresponding global mean, and masked non-brain tissue. Noteworthily, care was taken to minimize smoothing induced by interpolation and that no overt volume smoothing was performed.

The fMRI signal contains many sources of variability, including artifactual and non-neuronal signals, that make identifying the underlying neuronal activity difficult. Using a combination of independent component analysis (ICA) and classification techniques, HCP functional data were automatically denoised using FMRIB’s ICA-based X-noiseifier (Salimi-Khorshidi et al., 2014). Briefly, ICA was performed on each functional dataset independently and characteristics of each component, such as spatial localization and power in high frequencies, were evaluated by a classifier to determine if a given component was related to neuronal activity or artifact. The time-series corresponding to artifactual components were then regressed out of the data, providing a “cleaned”, denoised dataset for further investigation.

Using the minimally pre-processed, denoised resting-state datasets for each participant, the “global signal” was removed using FSL’s *fsl_glm* (Jenkinson et al., 2012) interface in NiPype (Gorgolewski et al., 2011). The “global signal”, although controversial in the domain of resting-state analyses, was removed under the premise that it performed better than other commonly used motion-correction strategies at removing motion-related artifacts in the HCP resting-state data (Burgess et al., 2016). The resulting data set was then smoothed with a FWHM kernel of 6-mm using FSL’s *susaan* interface in NyPipe. For each participant, the average time series for each ROI was extracted and a whole-brain correlation map was calculated and averaged across runs for a single participant for every ROI. The average correlation maps for each participant were transformed to Z-scores using Fisher’s r-to-z transformation. A group-level analysis was then performed to derive a rsFC map for each ROI using FSL’s *randomise* interface (Winkler et al., 2014) in NiPype. Images were thresholded non-parametrically using GRF-theory-based maximum height thresholding with a (voxel FWE-corrected) significance threshold of *P* < 0.001 (Worsley, 2001)), such that more spatially specific connectivity maps could be derived when using such a highly powered study (Woo et al., 2014).

### Task-Based Functional Connectivity: Meta-Analytic Co-Activation Modeling (MACM)

Leveraging reported coordinates from task-based fMRI studies, meta-analytic co-activation is a relatively new concept that identifies brain locations that are most likely to be co-activated with a given seed ROI across multiple task states. Differing from rsFC, MACM provides context about neural recruitment during goal-oriented behaviors. We therefore aimed to integrate these two complementary modalities by supplementing the rsFC maps with MACM maps for each ROI. To do so, we relied on the Neurosynth database (Yarkoni et al., 2011), which archives published stereotactic coordinates from over 14,000 fMRI studies and 150,000 brain locations. Neurosynth relies on an automated coordinate extraction tool to “scrape” each available fMRI study for reported coordinates. Due to the nature of this automated process, fMRI studies reporting results of multiple experimental contrasts as separate sets of coordinates are amalgamated into a single set of coordinates; in addition, “activation” and “de-activation” coordinates are not distinctly characterized. However, while this inherent “noise” may limit interpretational abilities, the power over manually curated datasets outweighs the potential confounds of bi-directional or mixed-contrast effects.

To generate a MACM map for each ROI, we utilized NiMARE (Salo et al., 2022) to search the Neurosynth database for all studies reporting at least one peak within the defined ROI mask. Neurosynth tools implement the multilevel kernel density analysis (MKDA) algorithm for performing meta-analyses based on a subset of studies, such as that described here. However, we opted to use the ALE algorithm as implemented in NiMARE given its optimal performance in replicating image-based meta- and mega-analyses (Salimi-Khorshidi et al., 2009). The ALE algorithm requires sample size information, or the number of subjects, that contributed to a given experimental contrast to generate a smoothing kernel. However, Neurosynth is not able to capture sample size (which could also vary across experimental contrasts within a study). Thus, we utilized a smoothing kernel with a FWHM of 15-mm, which has been shown to yield results with strong correspondence for image-based meta- and mega-analyses (SalimiKhorshidi et al., 2009). The ALE algorithm was applied to the set of studies reporting activation within the boundaries of each ROI. Once ALE maps were generated for each ROI, as described above, voxel-FWE correction (*P* < 0.001) was performed to reflect the statistical thresholding approach used for rsFC maps.

### Functional Decoding: Generalized Correspondence Latent Dirichlet Allocation (GC-LDA)

We sought to infer what mental processes were most likely linked with brain regions identified in our ALE, MACM, and rsFC analyses. To do so, we utilized generalized correspondence latent Dirichlet allocation (GC-LDA) functional decoding methods in NiMARE applied to the resulting unthresholded ALE, rsFC, and MACM maps. This type of decoding provides an approach to infer mental processes associated with neuroimaging spatial patterns. GC-LDA utilizes probabilistic Bayesian statistics that learns latent topics from a large database of papers (e.g., NeuroSynth) (Rubin et al., 2017). From the database, each topic found is treated as a probability distribution and creates a spatial distribution in MNI space across voxels from the maps entered into the decoding algorithm. The “topics” encompass terms and associated brain regions that co-occur in the literature from a literature database. We set our model to 200 topics. We report 10 terms corresponding to the highest weights associated with our ALE, rsFC, and MACM results.

## Results

### Literature Search and Study Criteria

The literature search yielded a total of 85 articles using the above-described search terms. **Figure 2** provides a PRISMA diagram, which details the review and filtering of those 85 studies. In the first round of review, records (i.e., titles and abstracts) were screened to exclude 18 studies that corresponded to non-human or non-English studies, reviews, or studies reporting white matter differences or differences among children or adolescents. Then, we examined the full-text articles to assess additional study criteria; 44 additional studies were excluded as being not eligible for the current meta-analysis.

**Figure 2.**
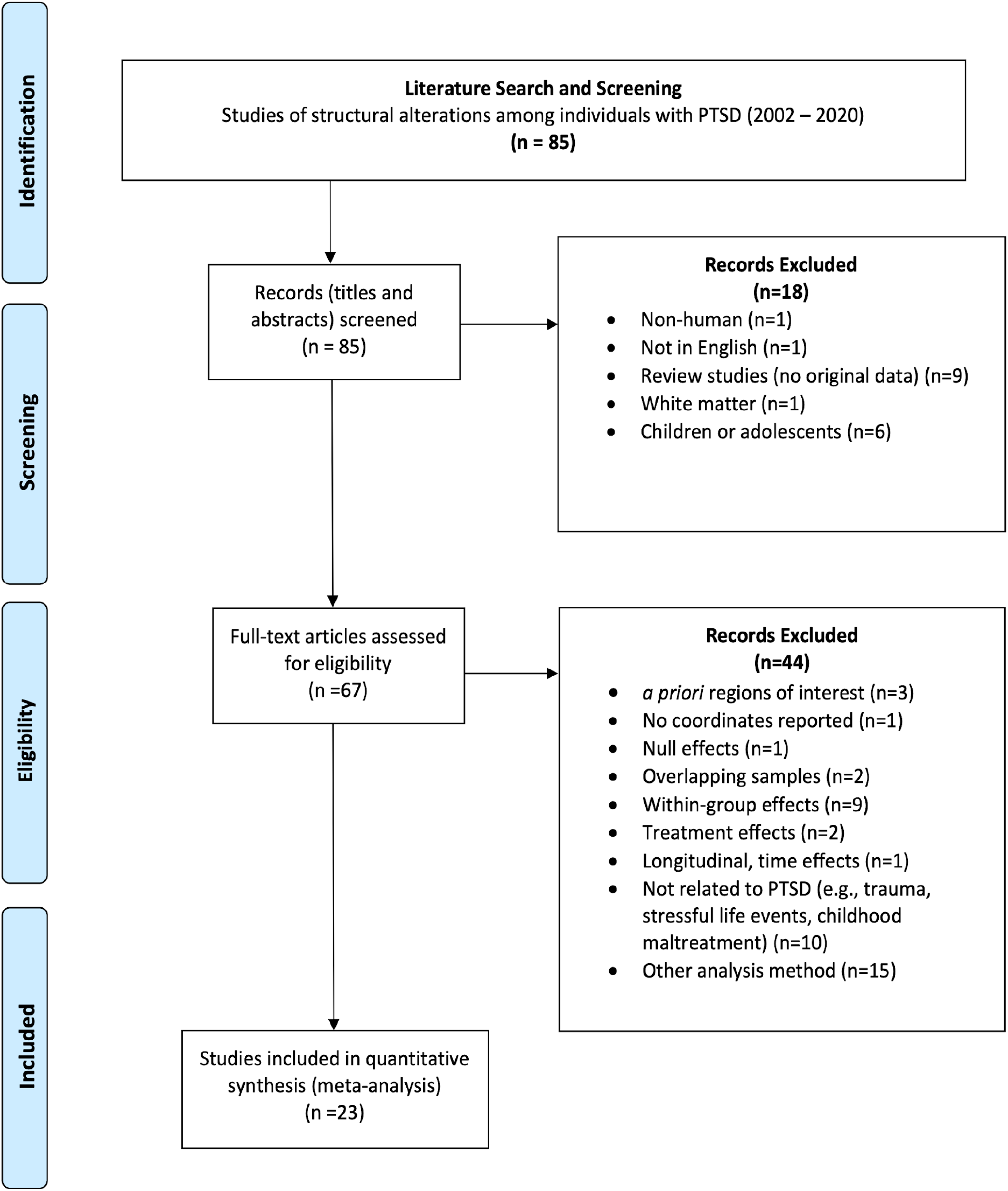
PRISMA Diagram. PRISMA flow chart detailing the literature search and selection criteria of studies included in the meta-analysis.

The final set of included studies consisted of 23 publications. Within these publications, gray matter structural alterations were assessed by comparing whole-brain VBM results among individuals with and without PTSD, reported as 3D coordinates in MNI or Talairach space. Control comparison groups included individuals who had experienced trauma but did not develop PTSD and individuals who had not experienced trauma. Nineteen publications included trauma-exposed controls (TC), while ten publications included healthy, non-trauma-exposed controls (HC). Altogether, this set of 23 studies collectively examined 476 individuals with PTSD and 892 individuals without PTSD, which included 288 TC and 633 HC. With respect to the type of structural alterations observed, studies reported multiple different VBM metrics. Seventeen publications reported group differences in gray matter volume (GMV), seven publications reported differences in gray matter density (GMD), and one reported gray matter concentration (GMC). Collectively, we refer to all of these metrics as gray matter (GM) differences among individuals with and without PTSD. Additional details on the demography of participant groups and study design are provided in **Supplementary Table 1** located in this project’s GitHub repository (https://github.com/NBCLab/meta-analysis_ptsd).

Within this final set of 23 publications, multiple contrasts of interest were reported. 25 contrasts reported GM *decreases* in PTSD vs non-PTSD for a total of 159 foci; this included 16 contrasts for PTSD vs. TC (82 foci) and 9 contrasts for PTSD vs. HC (77 foci). Conversely, 6 contrasts reported GM *increases* in PTSD vs. non-PTSD for a total of 20 foci, including 3 for PTSD (9 foci) vs. TC and 2 contrasts for PTSD vs. HC (9 foci).

### Anatomical Likelihood Estimation (ALE)

Using NiMARE v.0.0.3 (Salo et al., 2022), ALE meta-analysis was performed to assess convergence for the 25 contrasts from 22 publications of GM *decreases* among individuals with and without PTSD (i.e., non-PTSD > PTSD); a complete listing is provided in **Table 1**. Neuroimaging simulations indicate that a minimum of 20 contrasts are necessary for a well-powered coordinate-based meta-analysis (Eickhoff et al., 2016). Thus, we were unable to assess the 6 contrasts of GM *increases* (i.e., PTSD > non-PTSD) given insufficient power. With respect to GM decreases, we observed a single cluster of convergence located in the mPFC (x=0, y=46, z=10; BA 32) (**Figure 3**; *P* < 0.001, FWE-corrected *P* < 0.05). Given these results, we performed additional ALE meta-analyses for the PTSD vs. TC and PTSD vs. HC contrasts (i.e., GM increases and decreases) to determine if the use of different comparison groups potentially contributed additional heterogeneity, limiting assessment of convergence. However, we observed null results for these additional contrasts as well, likely in part due to the underpowered samples (Eickhoff et al., 2016).

**Table 1.** Studies Included in ALE Meta-Analysis. 25 contrasts from 22 publications reported GM *decreases* among individuals with and without PTSD (i.e., non-PTSD > PTSD). Sample sizes are provided for the total number of participants (N) (i.e., PTSD and non-PTSD), as well as the sample sizes for the PTSD groups (n).

|  | Citation | Sample Size | Contrasts |
| --- | --- | --- | --- |
| 1 | Bossini et al., 2017 | Total N=38; PTSD n=19 | Healthy Controls > PTSD |
| 2 | Chao et al., 2012 | Total N=41; PTSD n=21 | Non-PTSD > PTSD |
| 3 | Chen et al., 2009 | Total N=24; PTSD n=12 | Controls > PTSD |
| 4 | Chen et al., 2012 | Total N=20; PTSD n=10 | Controls > Recent Onset PTSD |
| 5 | Cheng et al., 2015 | Total N=60; PTSD n=30 | Healthy Controls > PTSD |
| 6 | Corbo et al., 2005 | Total N=28; PTSD n=14 | Healthy Controls > PTSD |
| 7 | Eckart et al., 2011 | Total N=33; PTSD n=20 | Non-Trauma Controls > PTSD |
| 8 | Felmington et al., 2009 | Total N=38; PTSD n=21 | Controls > PTSD |
| 9 | Hakamata et al., 2007 | Total N=184; PTSD n=14 | Non-PTSD > PTSD; Trauma Exposed > PTSD |
| 10 | Herringa et al., 2012 | Total N=28; PTSD n=13 | Trauma Exposed > PTSD |
| 11 | Kasai et al., 2008 | Total N=41; PTSD n=18 | Combat-Exposed Non-PTSD > PTSD |
| 12 | Kroes et al., 2011 | Total N=53; PTSD n=24 | Controls > PTSD |
| 13 | Li et al., 2006 | Total N=24; PTSD n=12 | Controls > PTSD |
| 14 | Nardo et al., 2010 | Total N=43; PTSD n=21 | Non-PTSD > PTSD |
| 15 | O'Doherty et al., 2017 | Total N=75; PTSD n=25 | Healthy Controls > PTSD; Trauma Exposed > PTSD |
| 16 | Qi et al., 2020 | Total N=220; PTSD n=57 | Trauma Exposed > PTSD |
| 17 | Rocha-Rego et al., 2012 | Total N=32; PTSD n=16 | Trauma Exposed Controls > PTSD |
| 18 | Sui et al., 2010 | Total N=31; PTSD n=11 | Healthy Controls > PTSD; Trauma Exposed > PTSD |
| 19 | Tavanti et al., 2012 | Total N=50; PTSD n=25 | Healthy Controls > PTSD |
| 20 | Yamasue et al., 2003 | Total N=25; PTSD n=9 | Non-PTSD > PTSD |
| 21 | Zhang et al., 2011 | Total N=20; PTSD n=10 | Trauma-Exposed > PTSD |
| 22 | Zhang et al., 2018 | Total N=39; PTSD n=14 | Non-PTSD > PTSD |

**Figure 3.**
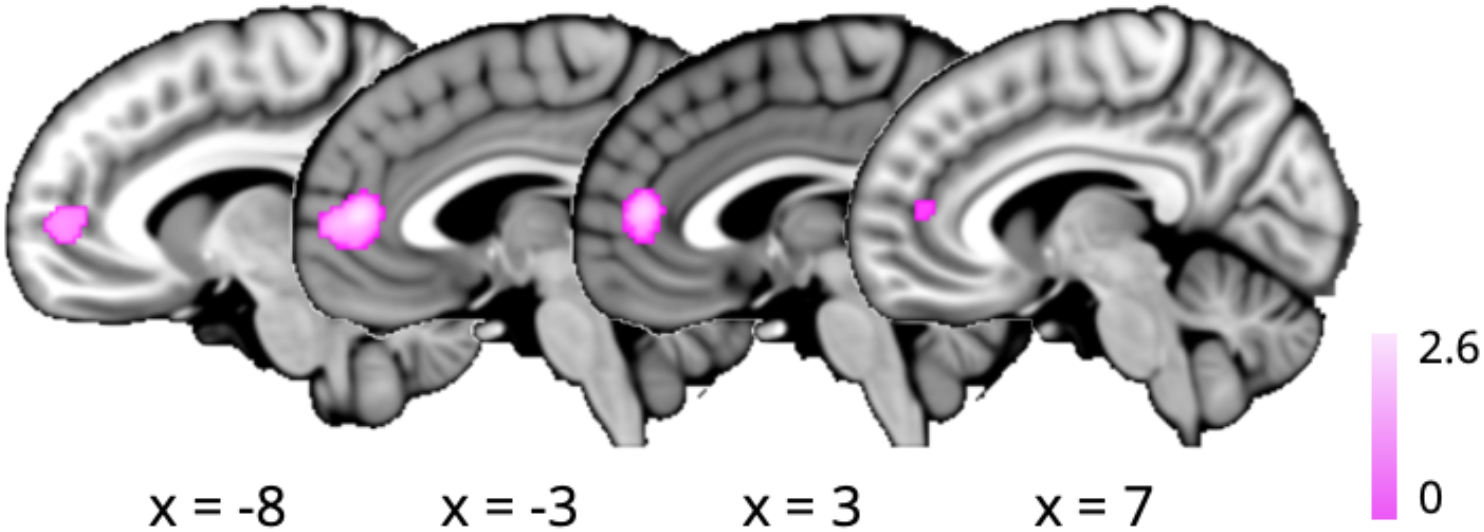
ALE Results for non-PTSD > PTSD. Sagittal brain slices illustrating convergent structural alterations associated with PTSD as determined by an ALE meta-analysis of GM reductions (*P* < 0.001, FWEcorrected *P* < 0.05).

### Functional Profiles of Structurally Altered Regions Associated with PTSD

We next investigated the functional connectivity of the mPFC cluster identified above showing convergent gray matter reductions among individuals with PTSD. To this end, we analyzed task-free rsFC and task-based MACM. First, we generated a rsFC map using the ALE-derived mPFC cluster as a seed region. The resultant rsFC map revealed rsFC with the superior frontal gyrus, medial frontal gyrus, inferior frontal gyrus, ACC, thalamus, posterior cingulate (PCC), superior temporal gyrus, medial temporal gyrus, precuneus, cuneus, and parahippocampus. Next, to further examine functionally coupled regions with the mPFC seed, we generated a MACM map using the Neurosynth database which demonstrated task-based coactivations with a similar pattern as the rsFC map. The locations of rsFC and MACM results are provided in **Table 2. Figure 4** illustrates the rsFC (blue) and MACM (red) results, with overlapping regions, indicating a consensus between rsFC and MACM (pink), revealed in the ACC, medial prefrontal gyrus, middle temporal gyrus, insula, inferior parietal lobe, thalamus, precuneus, parahippocampus, insula, and PCC regions (**Table 3**). This consensus pattern of regions suggests that PTSD-related gray matter loss in the mPFC may have implications for the optimal functioning of a widespread brain network.

**Table 2.**
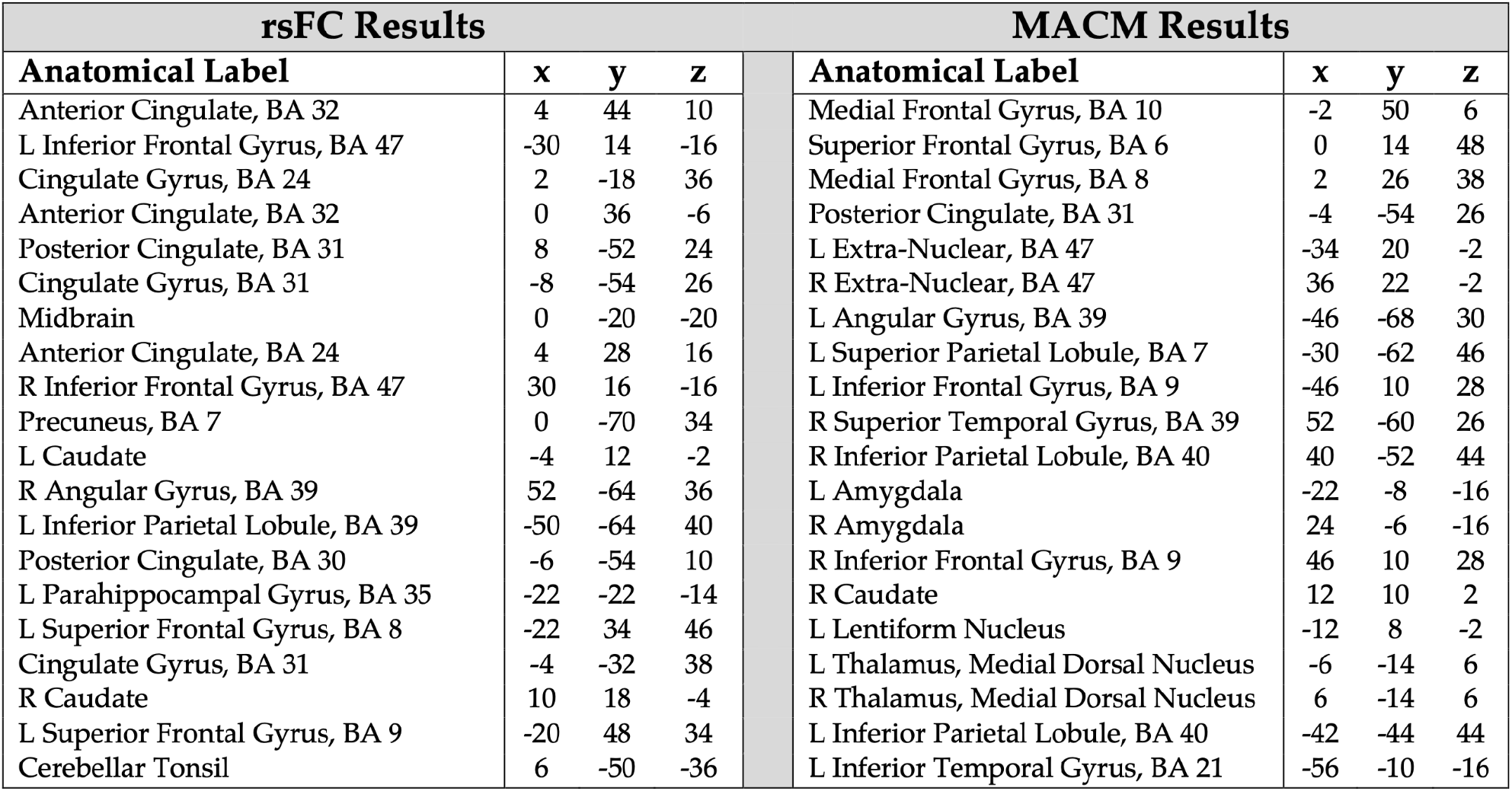
rsFC and MACM Results. Coordinate locations of the rsFC and MACM results, including the anatomical label and MNI coordinates of local maxima. Negative *x* values indicate the left (L) hemisphere and positive *x* values indicate the right (R) hemisphere.

**Table 3.** Consensus between rsFC and MACM Results. Coordinate locations of the consensus between rsFC and MACM results, including the anatomical label and MNI coordinates of local maxima. Negative *x* values indicate the left (L) hemisphere and positive *x* values indicate the right (R) hemisphere.

| rsFC + MACM Consensus |  |  |  |
| --- | --- | --- | --- |
| Anatomical Label | *x* | *y* | *z* |
| Medial Frontal Gyrus, BA 10 | -2 | 50 | 6 |
| Medial Frontal Gyrus, BA 8 | 2 | 26 | 38 |
| Posterior Cingulate, BA 31 | -4 | -54 | 26 |
| L Angular Gyrus, BA 39 | -46 | -68 | 30 |
| R Superior Temporal Gyrus, BA 39 | 52 | -60 | 26 |
| L Inferior Frontal Gyrus, BA 47 | -32 | 18 | -6 |
| R Inferior Frontal Gyrus, BA 47 | 38 | 20 | -8 |
| L Parahippocampal Gyrus, BA 28 | -24 | -16 | -18 |
| L Lentiform Nucleus, Putamen | -12 | 10 | -4 |
| R Caudate | 10 | 10 | -2 |
| L Thalamus, Medial Dorsal Nucleus | -4 | -14 | 6 |
| R Thalamus, Medial Dorsal Nucleus | 6 | -14 | 8 |
| L Parahippocampal Gyrus, BA 34 | -20 | 2 | -12 |
| R Hippocampus | 26 | -14 | -20 |
| L Inferior Temporal Gyrus, BA 21 | -56 | -10 | -16 |
| L Parahippocampal Gyrus, BA 28 | -16 | -4 | -14 |

**Figure 4.**
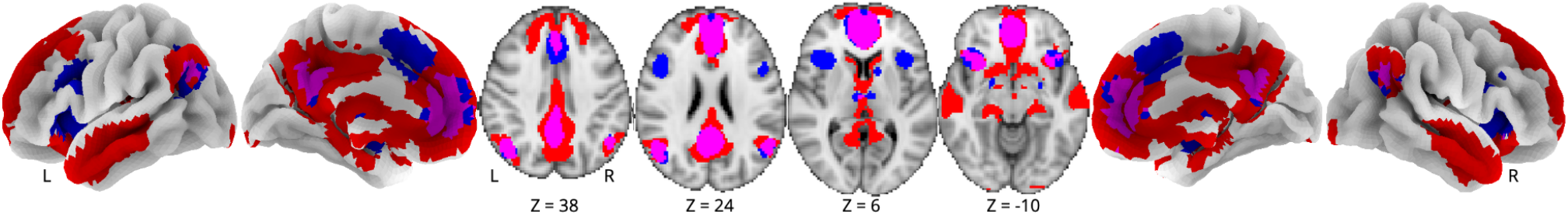
rsFC and MACM Results. rsFC (blue) and MACM (red) results; common areas (pink) indicate consensus between connectivity approaches. Images are thresholded at voxel-wise FWE *P* < 0.001.

### Functional Decoding: Generalized Correspondence Latent Dirichlet Allocation (GC-LDA)

Lastly, we performed functional decoding of the structural ALE, rsFC, and MACM maps to provide insight into the behavioral functions putatively associated with the observed functional connectivity patterns. Functional decoding was conducted using a GC-LDA analysis (Rubin et al., 2017). Because GCLDA does not provide correlational or statistical rankings, the top 10 unique terms computed from the GC-LDA analysis were taken into consideration separately for the structural ALE, rsFC, and MACM maps. The decoding terms with the top 10 weights from the GC-LDA analysis for the structural ALE map were: *visual, emotional, memory, novel, reward, motor, self, faces, learning, and face* (**Table 4a**). The decoding terms with the top 10 weights from the GC-LDA analysis for the rsFC map were: *default, default mode network, intrinsic, scale, self, person, reward, bias, judgements, and contexts* (**Table 4b**). Topographically speaking, the rsFC results resembled regions of combined default mode (Greicius et al., 2003; Raichle, 2015) and salience networks (Seeley et al., 2007; Menon and Uddin, 2010), and the functional decoding outcomes suggested that the rsFC results were associated with self-referential, intrinsic, and reward processes. Next, we examined MACM-based decoding results. The decoding terms with the top 10 weights from the GC-LDA analysis for the MACM map were: *visual, motor, emotional, memory, attention, auditory, reward, spatial, schizophrenia, and language* (**Table 4c**). Topographically speaking, the MACM results also resembled regions of the default mode (Greicius et al., 2003; Raichle, 2015) as well as the frontoparietal central executive network (Dosenbach et al., 2007; Seeley et al., 2007), and the functional decoding outcomes suggested association with executive emotional and memory processes. A summary of the decoding analyses for all three sets of images is shown as a radar plot in **Figure 5**.

**Table 4.** Functional Decoding Results. Functional decoding results for (a) ALE structural meta-analysis, (b) rsFC, and (c) MACM results as described by Neurosynth terms. Rankings display weighted terms listed from highest (1) to lowest (10).

| (a) ALE |  |  | (b) rsFC |  |  | (c) MACM |  |  |
| --- | --- | --- | --- | --- | --- | --- | --- | --- |
| Rank | Term | Weight | Rank | Term | Weight | Rank | Term | Weight |
| 1 | visual | 1.886 | 1 | default | 11.234 | 1 | visual | 5900.643 |
| 2 | emotional | 0.919 | 2 | default mode network | 9.225 | 2 | motor | 3839.578 |
| 3 | memory | 0.845 | 3 | intrinsic | 7.494 | 3 | emotional | 3665.765 |
| 4 | novel | 0.616 | 4 | scale | 6.236 | 4 | memory | 3476.688 |
| 5 | reward | 0.576 | 5 | self | 5.081 | 5 | attention | 2931.357 |
| 6 | motor | 0.521 | 6 | person | 4.977 | 6 | auditory | 2267.840 |
| 7 | self | 0.509 | 7 | reward | 4.780 | 7 | reward | 2107.441 |
| 8 | faces | 0.472 | 8 | bias | 4.568 | 8 | spatial | 2072.742 |
| 9 | learning | 0.467 | 9 | judgements | 4.279 | 9 | schizophrenia | 2070.157 |
| 10 | face | 0.450 | 10 | contexts | 4.271 | 10 | language | 2057.731 |

**Figure 5.**
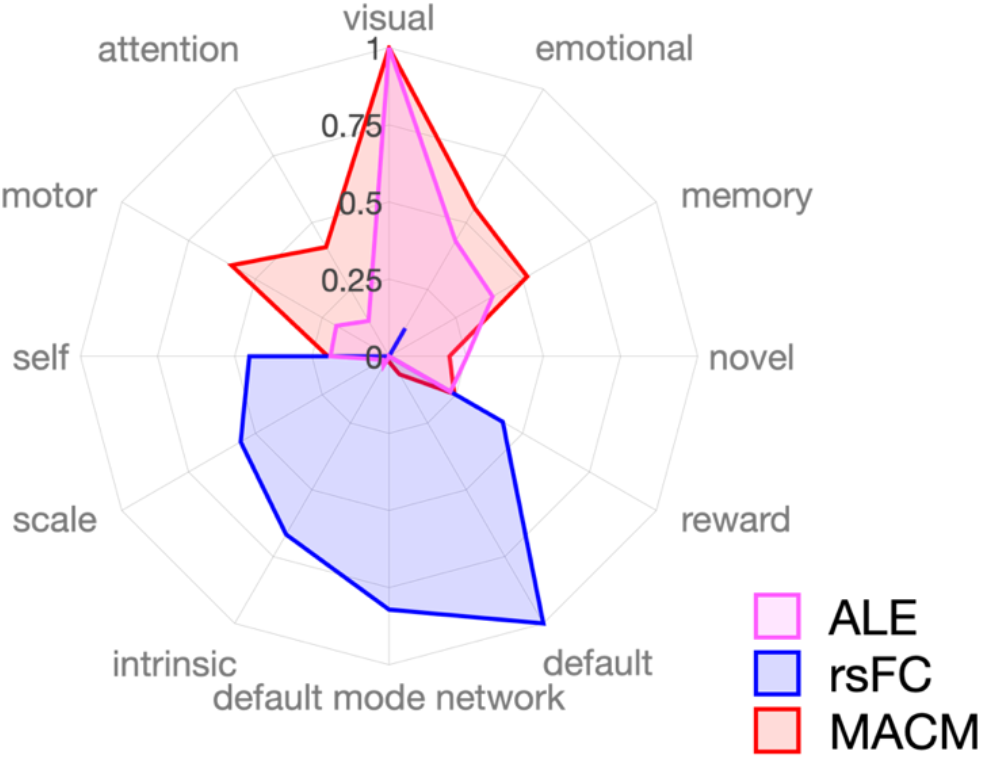
Functional Decoding Results. Functional decoding results for the ALE structural meta-analysis (pink), rsFC (blue), and MACM (red) results as described by Neurosynth terms. Radar plots display the top five terms across all three decoding analyses. The scale of the weights depends on both the GC-LDA model weights and the input values (Rubin et al., 2017); thus, the scale is arbitrary and has been normalized here to facilitate visualization.

## Discussion

The overall objective of this study was to investigate convergent alterations in brain structure among individuals with PTSD using emergent meta-analytic techniques. Further, we sought to extend the literature and assess potential functional consequences associated with observed structural alterations in PTSD by applying complementary rsFC and MACM analytic techniques. The current meta-analysis of 23 VBM studies evaluating GM volume alterations among PTSD versus non-PTSD groups identified a single node of convergent gray matter loss in the mPFC. GC-LDA-based functional decoding of this cluster was linked to Neurosynth terms of *visual, emotional, memory, novel, reward, motor, self, faces, learning, and face*. Follow-up ALE analyses exploring GM reductions in PTSD vs. HC (non-traumatized controls) and PTSD vs. TC (trauma-exposed controls not diagnosed with PTSD) yielded null findings likely due to insufficient power (Eickhoff et al., 2016). Subsequent analyses of the ALE-derived mPFC cluster were conducted to assess task-free (rsFC) and task-dependent (MACM) functional connectivity, identifying a consistent and widespread functional network implicated in PTSD. These results indicate that structural alterations in the mPFC among individuals with PTSD are possibly linked to disruptions across a larger frontoparietal network that includes the medial, superior, and inferior frontal gyri, PCC, parahippocampal gyri, angular gyri, superior temporal gyrus, thalamus, caudate, and lentiform nucleus. Functional decoding of rsFC and MACM results indicates substantive term overlap with the mPFC ALE results, with additional network-related terms (e.g., *default, default mode network, and intrinsic*).

### Structural Alterations and Dysfunction in PTSD

Our current findings suggest the mPFC appears as the most consistently reported brain region across VBM neuroimaging studies exploring the impact of PTSD on brain structure. Previous meta-analyses have identified GM reductions in the mPFC, hippocampus, fusiform gyrus, and lingual gyrus; however, not all of these regions were consistently observed across all meta-analyses (Kühn and Gallinat, 2013; Li et al., 2014; Meng et al., 2014; Bromis et al., 2018; Klaming et al., 2019; Serra-Blasco et al., 2021). Beyond the mPFC, we did not observe additional convergent GM reductions, indicating that prior findings in these other regions were not replicated. Across the PTSD literature, there is a high degree of variability associated with participant trauma exposure, length of diagnosis of PTSD, medication use, and comorbidity. Inconsistencies between our findings and previous meta-analytic results could be due to conceptual and methodological differences across the earlier studies, such as the scope of the research question exploring the neurobiology of PTSD, and the subsequent differences in inclusion/exclusion criteria that resulted in different sets of included studies. Comparison of the included studies in this and prior VBM meta-analyses of PTSD indicated varying degrees of overlap, including (from earliest to most recent meta-analyses): 7 of 9 included studies (Kühn and Gallinat, 2013), 14 of 17 included studies (Li et al., 2014), 15 of 20 included studies (Meng et al., 2014), 7 of 13 included studies (Bromis et al., 2018), 7 of 8 included studies (Klaming et al., 2019), and 10 out of 12 included studies (Serra-Blasco et al., 2021).

Beyond selection of included studies, the meta-analytic approach may contribute to the source of variability across results. Previous meta-analyses used either the ALE approach (Kühn and Gallinat, 2013; Li et al., 2014) or signed differential mapping (Meng et al., 2014; Bromis et al., 2018; Klaming et al., 2019; Serra-Blasco et al., 2021). Consistent with the present results, the meta-analyses by Meng et al. (2014) and Klaming et al. (2019) also yielded a single cluster in mPFC, which used the SDM method while our current results used the ALE approach. However, of all prior meta-analyses, only the study by Meng et al. (2014) meets the current threshold of a minimum of 20 contrasts for a well-powered coordinate-based meta-analysis (Eickhoff et al., 2016). After reviewing the above prior meta-analytic work in comparison to our current results, we conclude that extensive heterogeneity in the PTSD literature, combined with varying meta-analytic inclusive/exclusion criteria, likely contributed to differences between our results and prior meta-analytic findings. To our knowledge, the current meta-analysis of 25 contrasts represents the largest PTSD meta-analysis of structural findings to date, with prior meta-analytic work examining 8-20 included studies. We observed that the mPFC is robustly associated with structural alterations in PTSD; however, it is important to consider how the mPFC is integrated within existing neurocircuitry models associated with PTSD symptomology.

Traditional neurocircuitry models of PTSD utilize a fear-conditioning framework, emphasizing hyperreactivity of the amygdala in response to fear-related stimuli and dysfunction between the mPFC and orbitofrontal cortex, as well as the hippocampus, in attention and top-down control during threat exposure (Rauch et al., 2006; Shin et al., 2006). However, limiting consideration of the psychopathology of PTSD to focus on a single brain region (i.e., the amygdala) emphasizes fear-related brain activity while minimizing brain circuitry implicated in the complex constellation of PTSD symptoms associated with response to trauma exposure, such as re-experiencing trauma, avoidance, negative mood, and numbing. These additional processes remain largely unexplained in original PTSD models. However, more recent neurocircuitry models build from this perspective, with increased emphasis on altered function of the mPFC, its role in contextualization, and how context processing is core to the constellation of PTSD symptoms (Liberzon and Garfinkel, 2009; Liberzon and Abelson, 2016). While our results indicated convergent structural alterations in the mPFC, we did not observe similar convergence in the amygdala or other regions that have been implicated in prior neurocircuitry models of PTSD (Hamner, 1999; Rauch et al., 2006; Shin et al., 2006; Koenigs and Grafman, 2009). However, our results are congruent with the expanded models of PTSD and we provide robust evidence in support of the mPFC as a critical node in PTSD neurocircuitry. Further, our functional decoding results provide additional support for the contextualization models of PTSD. Taken together, reduced GM in the mPFC among individuals diagnosed with PTSD supports the premise that these structural alterations may contribute to deficits in context processing and ultimately play a dominant role in contributing to behaviors related to the constellation of symptoms in PTSD (Liberzon and Garfinkel, 2009; Liberzon and Abelson, 2016).

### Functional Profiles of Structural Findings in PTSD: Support for the Tripartite Model of Psychopathology

rsFC and MACM analyses characterized mPFC functional connectivity as extending across widespread, whole-brain networks engaging frontoparietal and limbic regions. These rsFC and MACM results, in conjunction with functional decoding outcomes, identified a functional connectivity profile suggestive of spatial patterns associated with the default mode network (DMN) (Greicius et al., 2003; Raichle, 2015), salience mode network (SN) (Seeley et al., 2007; Menon and Uddin, 2010), and central executive network (CEN) (Dosenbach et al., 2007; Seeley et al., 2007). The DMN is a system of connected brain areas including the mPFC, PCC, inferior parietal, and temporal cortices that are often collectively observed as displaying anticorrelation with regions actively engaged during attention-demanding tasks. Areas of the DMN are thought to collectively contribute to mental processes associated with introspection and self-referential thought (Liberzon and Shulman, 2001; Greicius et al., 2003; Whitfield-Gabrieli and Ford, 2012). The SN consists of the dorsolateral ACC and bilateral insula and is involved in saliency detection and attentional processes (Seeley et al., 2007; Menon and Uddin, 2010). Finally, the CEN consists of the dorsolateral prefrontal and posterior parietal cortices and is typically involved in attentionally driven cognitive functions, including goal-directed behavior (Turner et al., 2019). These three networks are central to a neurobiological theory of psychopathology (Menon and Uddin, 2010; Menon, 2011; Goodkind et al., 2015). The application of the tripartite model to neurobiology models of psychiatric disorders define dysfunction within and between connectivity of the DMN, SN, and CEN networks and relates to a broad range psychiatric disorders (Sha et al., 2019), including PTSD (Patel et al., 2012; Nicholson et al., 2020). Overall, the current meta-analysis identified a functional profile of the mPFC associated with connectivity between the DMN, SN, and CEN, which broadly supports a network theory of PTSD (Koch et al., 2016; Akiki et al., 2017).

According to the tripartite model of brain function, the SN is thought to mediate activity between the DMN and CEN networks in order to orient to external stimuli or internal salient biological stimuli (Menon and Uddin, 2010; Sripada et al., 2012a; Koch et al., 2016). Altered inter- and intra-network functional connectivity between the DMN, SN, and CEN has previously been implicated in PTSD (Koch et al., 2016). Specifically, seed-based resting state studies identified decreased connectivity within the DMN and SN, yet increased connectivity between these two networks among PTSD patients (Sripada et al., 2012a; Tursich et al., 2015). Furthermore, other resting state studies on PTSD utilizing graph theory approaches (Lei et al., 2015) and independent component analysis (Zhang et al., 2015) replicated weakened connectivity within the DMN, SN, and CEN, yet heightened connectivity between the DMN and SN (Tursich et al., 2015; Holmes et al., 2018). Taken together, this literature suggests deficits in top-down control over heightened responses to threatening stimuli and abnormal regulation of orienting attention to threatening stimuli (Sripada et al., 2012a; Lei et al., 2015; Tursich et al., 2015; Zhang et al., 2015; Koch et al., 2016). Patterns from task-based studies reflect previous findings of weakened connectivity between the SN and DMN and heightened connectivity between the SN and CEN (Patel et al., 2012; Thome et al., 2014). In a study among individuals with recent trauma exposure, connectivity between the DMN, SN, and CEN was reported to be disrupted among participants who developed PTSD vs. those who do not (Qin, 2012; Liu et al., 2017), providing evidence of differential functional connectivity between PTSD patients and traumatized non-diagnosed individuals. Network dysfunction associated with the DMN, SN, and CEN is also evident in task-based studies, including cues containing trauma stimuli (Rabellino et al., 2015), eye gaze (Thome et al., 2014), and a broad range of behavioral paradigms (Patel et al., 2012). Aberrant connectivity between and within the DMN, SN, and CEN has also been associated with PTSD symptoms, such that heightened connectivity and activity of the DMN was associated with depersonalization/derealization, while weakened connectivity and activity of the CEN was associated with hyperarousal and hypervigilance (Akiki et al., 2017). Additionally, weakened inter-network connectivity between the SN and DMN has been found to be positively correlated with Clinician Administered PTSD Scale (CAPS) scores that measure PTSD symptom severity (Sripada et al., 2012b; Tursich et al., 2015). Moreover, Bluhm et al. (Bluhm et al., 2009) found weakened spontaneous activity in regions of the DMN; in addition, posterior cingulate connectivity was positively correlated with self-reported dissociated experiences among participants with PTSD. In sum, the literature on abnormal brain function associated with PTSD points to a pattern of results suggesting that symptoms are related to aberrant connectivity within and between the DMN, SN, and CEN. In a recent review of the neuroimaging literature on PTSD, Lanius et al. (Lanius et al., 2015) summarized this work to reflect that dysfunction in the DMN is associated with an altered sense of self, dysfunction in the SN is associated with hyperarousal and hypervigilance, and dysfunction in the CEN is associated with cognitive dysfunction, including memory and cognitive control deficits.

The results from the current meta-analysis provide a robust mPFC-centric model of PTSD that is aligned with the extant literature and compliments the tripartite model of psychopathology. The mPFC, a core region of the DMN (Greicius et al., 2003; Raichle, 2015), is often disrupted in individuals with PTSD (Rabellino et al., 2015; DiGangi et al., 2016). The results of the present meta-analysis suggest alterations in mPFC structure, and related function, may play a crucial role in the underlying neurobiology of PTSD. Dysfunction of the mPFC is thought to be associated with poorer regulation of contextualization of PTSD symptoms. Prior literature indicates weakened integration of the DMN and disrupted inter-network connectivity with the SN and CEN, representing aberrant dysfunction of these tripartite networks in the psychopathology of PTSD (Ross and Cisler, 2020). Most of the prior functional and structural work involved varying analytic approaches, examined heterogeneous populations, and utilized region of interest approaches or *a priori* hypotheses. The current application of advanced meta-analytic techniques allowed for a whole-brain assessment of structural alterations associated with PTSD and the associated functional profiles of the mPFC. Future work in PTSD should consider integrating network-based analytic approaches with an mPFC-centric tripartite model to investigate differences in neuropathology of PTSD subtypes (e.g., trauma experiences, duration of exposures), characterizing heterogeneous presentations of PTSD symptoms, and potential predispositional developmental effects among youth, adolescent, and adult populations.

### Limitations

Our study is limited by several considerations. First, the present meta-analysis is limited by the small number of studies included. The studies that met the standards of inclusion for this study were considered to reduce instances of variance and consider reliability of study findings (inclusion and exclusion criteria are shown in Fig. 2). By considering the inclusion of trauma-exposed controls, healthy controls, and individuals with PTSD, the number of participants across each group was somewhat unevenly distributed due to small sample sizes in the original studies. However, the current meta-analysis met the previously recommended standard of at least 20 experimental contrasts required to conduct a well-powered meta-analysis (Eickhoff et al., 2016). Second, much heterogeneity exists across the studies included in our meta-analysis. For example, many of the studies included diagnostic criteria for PTSD using different clinical measures and reported different instances of duration of PTSD (e.g., lifetime vs. first onset). Substantial variability was also present in the type of trauma and duration of exposure to trauma within the different groups for this study. Given these issues, we were unable to classify PTSD subtypes across the included studies and thus have reported results that relate to generalized PTSD. Many of the original studies were not able to clearly disentangle comorbidity of PTSD with other psychiatric disorders (e.g., depression, anxiety) or report instances of medication and drug abuse. Furthermore, studies relied on various neuroimaging acquisition and analysis methods, which likely introduced additional variability associated with methodological flexibility (Carp, 2012; BotvinikNezer et al., 2020). However, the goal of neuroimaging meta-analysis is to examine consensus despite such variability in the literature. With this in mind, we are confident that the mPFC is a significant brain region linked to GM reductions in PTSD, as well as a robust node of the DMN that plays an important role in toggling between the DMN, SN, and CEN. Future transdiagnostic and meta-analytic work is needed to identify similar and unique neurobiological mechanisms of PTSD in comparison to other related disorders, including complementary disease-decoding or structural covariance analysis, which would further advance clinical insight.

## Conclusions

The present study utilized coordinate-based meta-analytic techniques to determine that reduced mPFC GM is consistently found among individuals with PTSD. Complementary analyses of rsFC and MACM functional connectivity provided novel insight into how structural alterations may have potential functional consequences. Our results indicated that decreases in mPFC GM may be linked to widespread functional systems that are implicated in behavioral deficits and cluster symptomatology of PTSD. Specifically, consensus-based functional profiles, across task-free and task-based domains, emphasized brain regions associated with the tripartite model of psychiatric disorders where inter- and intra-network connectivity involving the DMN, SN, and CEN are core to PTSD dysfunction. Overall, these results may be important in providing a more comprehensive understanding of the neurobiological bases of PTSD, which is needed to understand the varying diagnosis, symptomatology, and treatment of PTSD, as well as enhanced targeting of treatment towards heterogeneous classification and symptom clusters of PTSD.

## Declarations

## Ethics Approval

This secondary data analysis was approved by the Institutional Review Board of Florida International University.

## Consent to Participate

Not applicable.

## Consent for Publication

Not applicable.

## Availability of Data and Materials

Data and materials are available in a GitHub repository (https://github.com/NBCLab/meta-analysis_ptsd), including the meta-analytic coordinate files, data analysis scripts (i.e., code), image-based results (i.e., ALE, rsFC, and MACM images), and functional decoding results. rsFC analyses used the Human Connectome Project’s (Van Essen et al., 2013) Young Adult Study S1200 Data Release (March 1, 2017), which is available at db.humanconnectome.org.

## Author Contributions

BP and IC collected and prepared data for meta-analysis. MCR and JEB analyzed data. MCR, JEB, LDHB, and TS contributed scripts and pipelines. BP, MRC, and ARL wrote the paper. All authors contributed to the revisions and approved the final version.

## Acknowledgments

The authors would like to thank the FIU Instructional & Research Computing Center (IRCC, http://ircc.fiu.edu) for providing the HPC and computing resources that contributed to the research results reported within this paper.

